# Analysis of the mechanism of Aldo-keto reductase dependent cis-platin resistance in HepG2 based on transcriptomic and NADH metabolic analysis

**DOI:** 10.1101/2021.04.29.441897

**Authors:** Tingting Sun, Xue Sun, Xin Wang, Rui Guo, Yuanhua Yu, Le Gao

## Abstract

Aldo-keto oxidoreductase (AKR) inhibitors could reverse several cancer cells’ resistance to Cis-platin, but their role in resistance remains unclear. Our RNA-seq results showed de novo NAD biosynthesis-related genes, and NAD(P)H-dependent oxidoreductases were significantly upregulated in Cis-platin-resistant HepG2 hepatic cancer cells (HepG2-RC cells) compared with HepG2 cells. Knockdown of AKR1Cs could increase Cis-platin sensitivity in HepG2-RC cells about two-fold. Interestingly, the AKR1C inhibitor meclofenamic acid could increase Cis-platin sensitivity of HepG2-RC cells about eight-fold, indicating that knockdown of AKR1Cs only partially reversed the resistance. Meanwhile, the amount of total NAD and the ratio of NADH/NAD+ were increased in HepG2-RC cells compared with HepG2 cells. The increased NADH could be explained as a directly operating antioxidant to scavenge radicals induced by Cis-platin. We report here that NADH, which is produced by NAD(P)H-dependent oxidoreductases, plays a key role in the AKR-associated Cis-platin resistance of HepG2 hepatic cancer cells.

## 1 Introduction

Aldo-keto reductases (AKRs) are NAD(P)H-dependent oxidoreductases, which reduce carbonyl substrates with NAD(P)H and are present in all three domains of life. Human AKRs are important in metabolic pathways such as steroid biosynthesis, alcohol oxidation, and xenobiotic elimination[1]. Some AKRs, such as AKR1B10[2] and AKR1Cs[3,4], are correlated with carcinogenesis. AKR1Cs have also been related to Cis-platin resistance in gastric carcinoma TSGH-S3 cells[5] and metastatic bladder cancer cells[6].

Cis-platin is a front-line chemotherapy agent for cancer treatment[7]. The AKR1C inhibitor flufenamic acid was found useful to reverse Cis-platin resistance of bladder cancer[6]. It was speculated that AKR1Cs could reduce some cytotoxic lipid peroxidative products from aldehydes[5]. Hepatic cancer cells with high expression levels of AKRs usually show a rather high tolerance to Cis-platin treatment. However, the target molecule of AKRs remains unknown. Here we report that NADH, a product of Aldo-keto oxidation-reduction, plays a key role in the Cis-platin resistance of HepG2 hepatic cancer cells.

## 2 Materials and Methods

### 2.1 Cell culture

Human hepatic HL-7702 cells and the hepatic cancer cell lines HepG2 and its Cis-platin-resistant strain HepG2-RC were purchased from Fusheng Biotechnology (Shanghai, China). HL-7702 cells were cultured at 5% CO2 in RPMI-1640 medium. HepG2 cells were cultured at 5% CO2 in DMEM medium. HepG2-RC cells were cultured at 5% CO2 in MEME medium with increasing Cis-platin concentration until 50% of cells died.

### 2.2 Quantitative real-time polymerase chain reaction

Total RNA isolation and first-strand cDNA synthesis from HL-7702, HepG2, and HepG2-RC cells were performed by a SuperReal Kit (Tiangen, Beijing, China). The primer sequences are listed in Table S1. Quantitative real-time polymerase chain reaction (qRT-PCR) was performed using ABI7900HTFast (Thermofisher, Waltham, MA). The data were normalized to the β-actin expression level and are expressed as the fold change relative to control (2^−ΔΔCt^).

### 2.3 Western blot

Cells were washed twice with cold PBS and lysed in a buffer containing 0.5% NP-40, 10 mM Tris-HCl (pH 7.4), 150 mM NaCl, 1 mM EDTA, 50 mM NaF, 1 mM PMSF, and 1 mM Na3VO4, and the lysate was clarified by centrifugation at 15,000 rpm for 10 min. The supernatants were then subjected to 8%–12% SDS-PAGE. Separated proteins were transferred to polyvinylidene difluoride membranes and blocked using Tris-buffered saline containing Tween-20 (TBS-T) with 5% skim milk at room temperature for 1 h. Membranes were incubated with the following primary antibodies: anti-AKR1C1 (Abnova, Taiwan, China), anti-AKR1C2 (Abcam, Cambridge, UK), anti-AKR1C3 (Abcam, Cambridge, UK), and AKR1C4 (Abnova, Taiwan, China) at 4°C overnight. Membranes were washed with TBS-T, followed by incubation with responding secondary antibodies. The membranes were washed three times in TBS-T, and the signal was developed using ECL (GE Healthcare, Little Chalfont, UK), followed by detection using an AzureC600 detection system (Azure Biosystems, Dublin, CA).

### 2.4 Measuring IC50 of AKR1C inhibitors

The activity of AKR1C3 was estimated by measuring the OD340 (NADH) during the conversion of glyceraldehyde to glycerol by the Vallee–Hoch method. Three inhibitors, medroxyprogesterone acetate (MPA), meclofenamic acid (MCFLA), and methyliasmonate (MLS), were applied to inhibit AKR1C3 activity. The IC50 of the inhibitors was calculated from the plots of AKR1C3 activity vs concentration of inhibitors. Each sample was measured in triplicate.

### 2.5 Reverse Cis-platin resistance

MPA, MCFLA, and MLS were applied to reverse Cis-platin resistance of HepG2-RC. The inhibitors at given concentrations (MPA: 0.31 mM, MCFLA: 0.12 mM, MLS: 0.13 mM) were co-incubated with HepG2-RC cells under a gradient concentration of Cis-platin. The Cis-platin IC50 of each inhibitor-treated HepG2-RC sample was then calculated from the MTT assay.

For the knockdown of AKR1Cs in HepG2-RC cells. Small interfering RNAs (siRNAs) targeting human AKR1C1-4 were synthesized by Qiagen (Valencia, CA). HepG2-RC cells (5×106 cells/well) were transfected with 50 nM of si-AKR1Cs or si-scramble as control (sequences are provided in Table S1) using HiPerfect transfection reagent (Qiagen, Valencia, CA). After 24 or 48 h transfection, cells were subjected to RT-PCR and immunoblotting to identify whether AKR1Cs were knocked down. The HepG2-RC cells in which knockdown of AKR1Cs was successful then underwent a Cis-platin IC50 test as described above.

### 2.6 RNA-sequencing and data analysis

Approximately 106 HepG2 and HepG2-RC cells were frozen on dry ice. RNA extraction, library preparation, RNA-seq, and bioinformatics analysis were performed at BGI (Shenzhen, China). Each set of cell samples was sequenced in three independent experiments. Image analysis, base-calling, and filtering based on fluorescence purity and output of filtered sequencing files were performed through the Illumina analysis pipeline.

The obtained raw reads of HepG2 and HepG2-RC cells were preprocessed by removing reads containing adapter sequences, reads containing poly-N, and low-quality reads. Q20 and Q30 were calculated. All six runs of HepG2 and HepG2-RC samples showed that at least 96% of reads was Q20, and at least 87% was Q30. All downstream analyses were based on high-quality clean reads. Gene function was annotated based on Gene Ontology (GO) and the Kyoto Encyclopedia of Genes and Genomes (KEGG). Genes with log2(fold change) > 1 and FPKM > 0.1 were selected as upregulated genes. Genes with log2(fold change) < −1 and FPKM > 0.1 were selected as downregulated genes. Finally, 1486 upregulated genes and 270 downregulated genes were identified.

### 2.7 Assay of NADH and NAD+ content

The amounts of NADH and NAD+ were measured by NAD+/NADH Assay Kit with WST-8 (Beyotime, Shanghai, China). Approximately 106 HL-7702, HepG2, and HepG2-RC cells were collected for NAD+ or NADH extraction. Each sample was divided into two equal parts. One part was used for measuring the total amount of NAD (NAD++NADH), and the other part was used for measuring the amount of NADH after 60°C heat treatment. In the assay, NAD+ was first converted to NADH by adding alcohol dehydrogenase and ethanol. NADH then reduced WST-8 to formazan, and the amount of NADH could be measured by monitoring the OD450 value. The ratio of NAD+/NADH was calculated by the formula *ratio* = ([*NAD*]*_total_* – [*NADH*])/([*NADH*]). Each sample was measured in three independent experiments.

### 2.8 Docking experiment

A protein docking model of AKR1C3 and its inhibitors was simulated by AutoDock 4.2.6 (The Scripps Research Institute, San Diego, CA). The crystal structure of AKR1C3 was derived from the RCSB protein data bank (ID: 4DBW). Ligands were initially drawn using ChemBioDraw Ultra 13.0 (PerkinElmer, Waltham, MA), and energy minimization was performed with ChemBio3D Ultra 13.0 using the MMFF94 force field (PerkinElmer, Waltham, CA). Optimized ligand candidates were saved in PDBQT format. The dimensions of the grid box were set at 110, 110, 85 (x, y, z), and the center of the box was placed on Tyr-55 in the A-chain. MPA, MCFLA, and MLS were designed and docked onto the AKR1C3 model with or without NADP+, and their binding energy was estimated from docking models.

## 3 Results

### 3.1 AKR1C3 was upregulated in HepG2-RC cells compared with HepG2 cells

To investigate the differential expression of AKR1Cs in HepG2 and HepG2-RC cells, we evaluated their mRNA and protein levels. According to the qRT-PCR results (Table 1), the mRNA levels of all AKR1C isoenzymes were higher in HepG2 cells compared with human hepatic HL-7702 cells. However, only AKR1C3 was upregulated in HepG2-RC cells compared with HepG2 cells (~50-fold), while the other isoenzymes showed decreased levels in HepG2-RC cells.

**Table.1.**
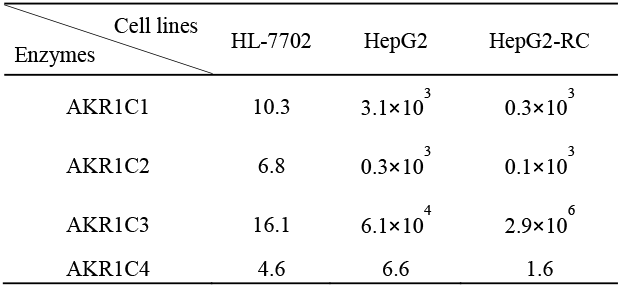
The mRNA amount (2-ΔCt) of AKR1Cs in different hepatic cell lines measured by qRT-PCR.

According to our Western blot results (Fig.1), AKR1C1, AKR1C2, and AKR1C4 levels were almost equal between HepG2 and HepG2-RC cells (1.2-fold differences). AKR1C3 levels in HepG2-RC cells were as much as 1.5-fold higher than in HepG2 cells, which was consistent with our qRT-PCR results.

**Fig. 1.**
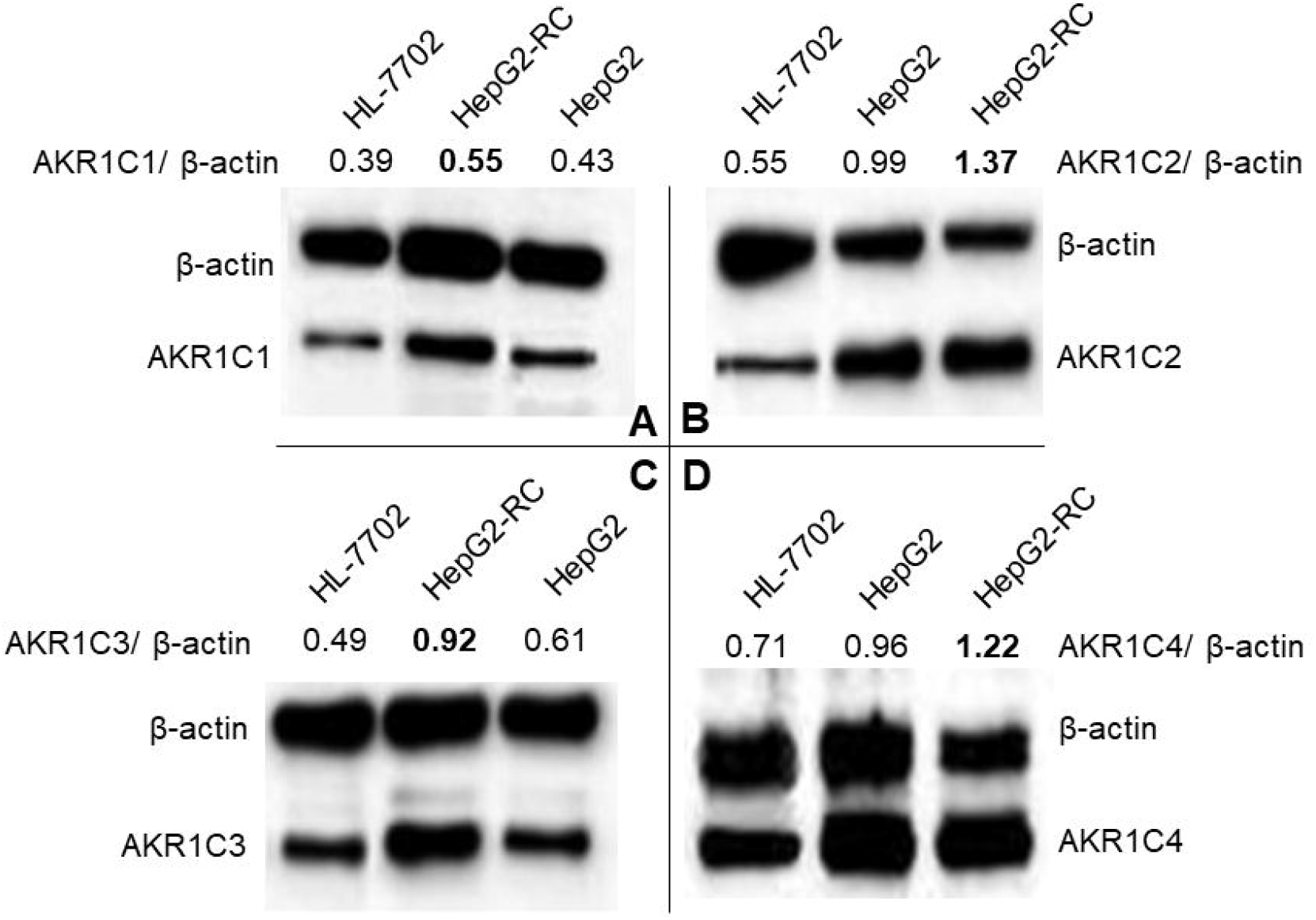
Differential expression of AKR1Cs in hepatic cell lines HL-7702, HepG2, and HepG2-RC, as measured with western blot. A, AKR1C1; B, AKR1C2; C, AKR1C3; D, AKR1C4. β-actin was selected as an internal reference. The signal intensity of each band was assessed with ImageJ software and divided by the value of β-actin. The ratios of AKR1Cs and β-actin are shown at the top of the figure.

### 3.2 AKR1C inhibitors could reverse Cis-platin resistance of HepG2-RC cells

Three AKR1C inhibitors, MPA, MCFLA, and MLS, were applied to reverse Cis-platin resistance in HepG2-RC cells. First, the IC50 value of each AKR1C3 inhibitor was determined (Fig. 2A). MPA showed the lowest IC50 value (2.1 μM). The IC50 values of the other inhibitors were 3.3 μM (MCFLA) and 16.3 μM (MLS). However, MCFLA caused the strongest increase in Cis-platin sensitivity (~8-fold). MPA and MLS increased Cis-platin sensitivity almost 2.5-fold and approximately 1.5-fold, respectively (Fig. 2B).

**Fig. 2.**
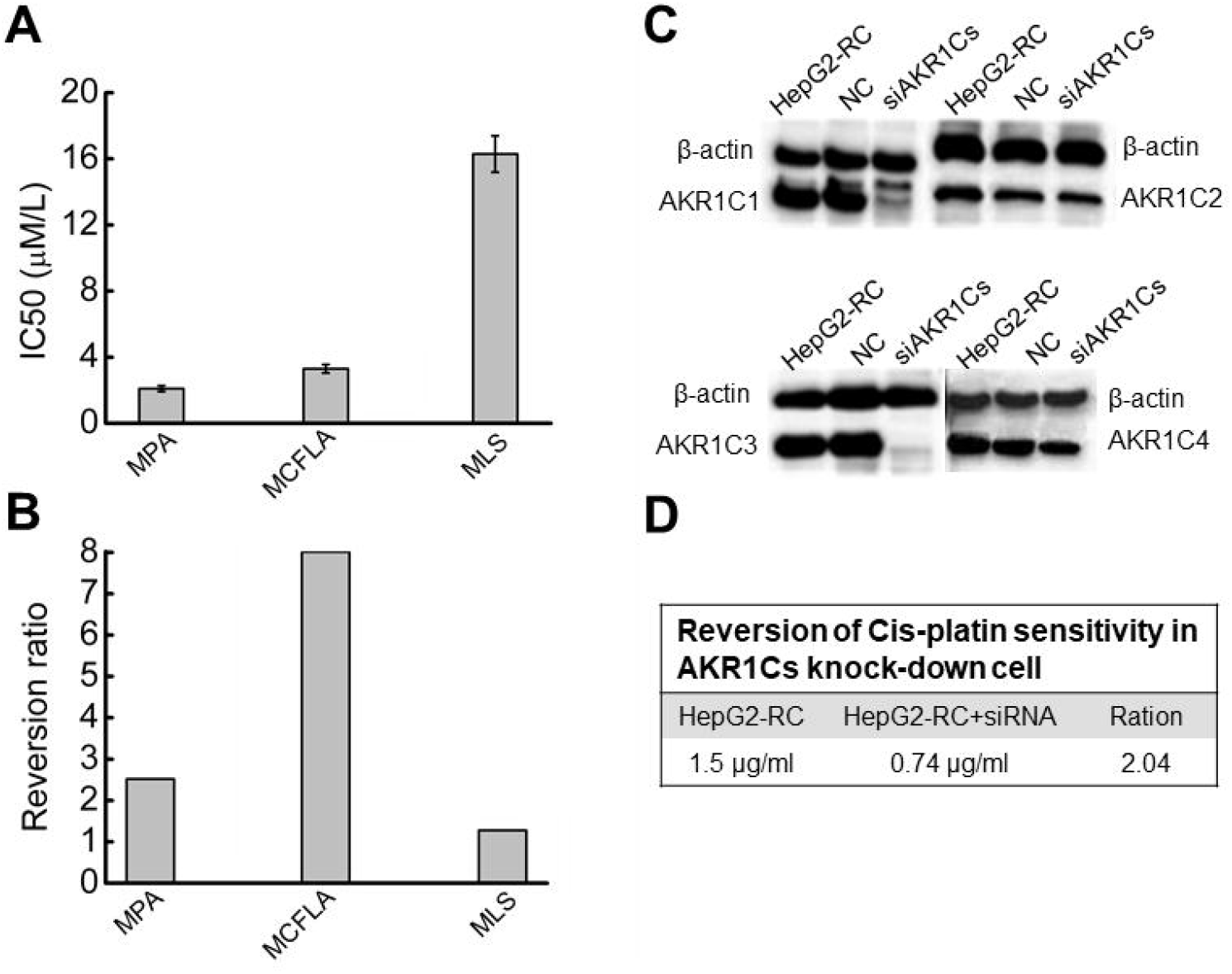
Reversion of Cis-platin sensitivity in HepG2-RC. A, IC50 value of three inhibitors acting on AKR1C3. MPA, medroxyprogesterone acetate; MCFLA, meclofenamic acid; MLS, methyliasmonate. B, Reversion ratio of Cis-platin sensitivity in HepG2-RC cells caused by three inhibitors. C, Knockdown of AKR1Cs by siRNA. Successful knockdown was confirmed with western blot. D, Reversion of Cis-platin sensitivity in AKR1C knockdown cells.

### 3.3 SiRNA of AKR1Cs could partially reverse resistance of HepG2-RC cells

Since AKR1C inhibitors could reverse Cis-platin resistance of HepG2-RC cells, RNAi knockdown experiments of all AKR1Cs were performed. Western blot results showed that AKR1C1 and AKR1C3 protein levels were strongly reduced, while these remained unchanged in control-siRNA HepG2-RC cells (Fig. 2C). These observations confirm that AKR1C1 and AKR1C3 were successfully knocked down in HepG2-RC cells.

Cis-platin resistance reversal experiments showed that the AKR1C knockdown HepG2-RC cells could tolerate two-fold lower Cis-platin concentrations than control HepG2-RC cells (Fig. 2D). The effects of AKR1C knockdown on Cis-platin resistance were equivalent to the effects of MPA (~2.5-fold reversal), but were much weaker than the effects of MCFLA (~8-fold reversal), indicating that knockdown of AKR1Cs could partially reverse Cis-platin resistance of HepG2-RC cells.

### 3.4 Most NAD(P)H-dependent reductase/oxidases were upregulated in HepG2-RC cells

According to our RNA-seq results, AKR1C levels were not greatly changed in HepG2-RC cells (log2(HEPG2-RC/HEPG2) < 2; Table 2), and only AKR1C3 showed a slight increase, while the other three showed a slight decrease. This tendency was consistent with the above qRT-PCR results. Among the 16 human AKR enzymes, four (AKR1B10, AKR1B15, AKR1D1, and AKR1B1) were upregulated about four-fold, while two (AKR1E2 and AKR1C4) were downregulated about twofold; the remaining nine enzymes showed almost no change.

**Table.2.** Regulation of AKR enzymes measured by RNA-seq.

| Gene | log2(HepG2-RC/HepG2) | Annotation |
| --- | --- | --- |
| AKR1B10 | 3.12 | All-trans-retinol + NADP+ ⇌ all-trans-retinal + NADPH |
| AKR1B15 | 2.63 | L-Arabitol:NADP + 1-oxidoreductase L-Arabitol + NADP+ ⇌ L-Arabinose + NADPH + H+ |
| AKR1D1 | 2.16 | 5β-androstane-3,17-dione:NADP + 4,5-oxidoreductase 5β-Androstane-3,17-dione + NADP+ ⇌ H+ + NADPH + Androstenedione |
| AKR1B1 | 2.14 | glycerol:NADP + oxidoreductase Glycerol + NADP+ ⇌ |
| AKR1C3 | 0.41 | D-Glyceraldehyde + NADPH + H+ |
| AKR6A5 | 0.21 | NADP+ + trans-1,2-dihydrobenzene-1,2-diol ⇌ catechol + H+ + NADPH |
| AKR6A9 | 0.18 |  |
| AKR7A3 | -0.02 | potassium voltage-gated channel subfamily A regulatory beta subunit 2 |
| AKR6A3 | -0.09 |  |
| AKR1C1 | -0.43 | potassium voltage-gated channel subfamily A regulatory beta subunit 3 |
| AKR1C2 | -0.44 | aflatoxin B1 + NAD(P)+ ⇌ aflatoxin B1-dialchol + NAD(P)H |
| AKR7A2 | -0.55 | potassium voltage-gated channel subfamily A regulatory beta subunit 1 |
| AKR1A1 | -0.68 |  |
| AKR7A4 | -0.94 | 17α,20α-dihydroxypregn-4-en-3-one + NAD(P)+ ⇌ |
| AKR1E2 | -1.03 | 17α-hydroxyprogesterone + H+ + NAD(P)H |
| | | 3α-hydroxysteroid + NADP+ ⇌ 3-oxosteroid + H+ + NADPH |
| AKR1C4 | -1.03 | 4-hydroxybutanoate + NADP+ ⇌ H+ + NADPH + succinate semialdehyde |
| | | allyl alcohol + NADP+ ⇌ acrolein + H+ + NADPH |
| | | aflatoxin B1 + NAD(P)+ ⇌ aflatoxin B1-dialchol + NAD(P)H |
| | | testosterone:NAD+ 17-oxidoreductase Testosterone + NAD+ ⇌ Androstenedione + NADH + H+ |
| | | Androsterone:NADP+ oxidoreductase Androsterone + NADP+ ⇌ 5α-Androstane-3,17-dione + NADPH + H+ |

Moreover, many other NAD(P)H-dependent reductase/oxidases were upregulated four- to eight-fold (Table 2). The strongest upregulation was observed for RFTN1 (log2(HEPG2-RC/HEPG2) = 6.55), which encodes raftlin, which bears NAD(P)H cytochrome-b5 reductase activity. Comparing RNA-seq results from HepG2 and HepG2-RC cells, 63 NAD(P)H-dependent reductase/oxidases were upregulated in HepG2-RC cells at least twofold, while only 23 were downregulated at least/approximately twofold. Moreover, 23 of those 63 upregulated genes had at least four-fold higher transcription in HepG2-RC cells compared with HepG2 cells, while only two of those 23 downregulated genes showed a reduction to less than 25% of HepG2 levels. In other words, even though no NAD(P)H-dependent reductase/oxidases were present in the top 10 upregulated genes (Tables S2 and S3), they were generally upregulated.

### 3.5 Ratio of NADH/NAD+ in HepG2-RC cells was higher than in HepG2 cells

As shown in Fig. 3, the amount of total NAD in HepG2 and HepG2-RC cells was approximately four-fold higher than in HL-7702 cells. Interestingly, the ratio of NADH/NAD+ in HepG2-RC cells was almost seven-fold higher than in HepG2 or HL-7702 cells. These results could be explained by the RNA-seq results that de novo NAD biosynthesis-related genes were upregulated in HepG2-RC cells (Table 3). Especially, TDO2, encoding tryptophan dioxygenase; KYNU, encoding kynureninase; and NMNAT2, encoding nicotinamide nucleotide adenylyl transferase, showed at least four-fold and at most 32-fold higher mRNA levels in HepG2-RC than in HepG2 cells. Moreover, almost all genes involved in NAD degradation did not show altered mRNA levels, except the ART family. All ART family genes were upregulated in HepG2-RC compared with HepG2; in particular, ART1 mRNA levels in HepG2-RC cells increased 32-fold compared with HepG2 cells. It has been reported that ART1 overexpression is closely related to some kinds of cancer[8–10]. In summary, the de novo NAD biosynthesis pathway was upregulated in HepG2-RC, and total NAD levels were increased in HepG2 and HepG2-RC cells compared with HL-7702 cells.

**Fig. 3.**
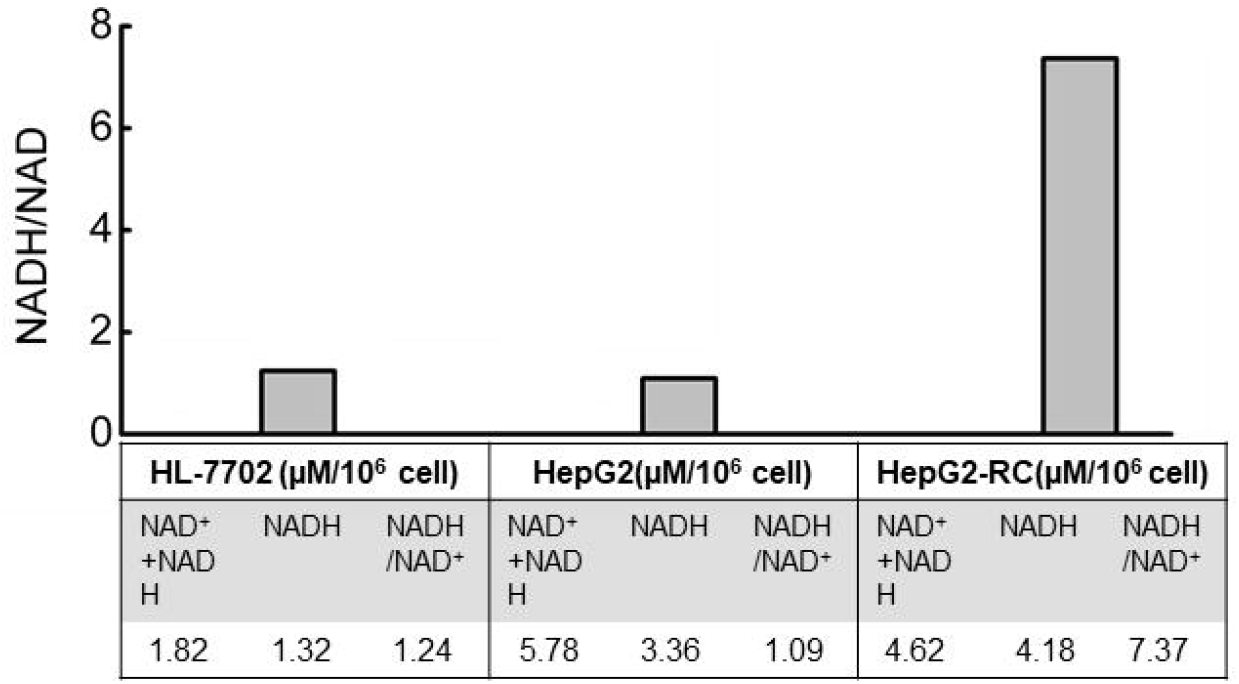
Ratio of NADH/NAD+ in HL-7702, HepG2, and HepG2-RC cells. The amounts of total (NAD++NADH) and NADH were measured with an NAD/NADH quantitation kit. The amount of NAD was calculated from these two values. Finally, a ratio of NADH vs NAD was obtained.

**Table.3.** Regulation of NAD metabolism related genes in HepG2 and HepG2-RC measured by RNA-seq.

| Gene | Log2(HepG2-RC/HepG2) | Annotation |
| --- | --- | --- |
| IDO1 | 0.41 | indoleamine 2,3-dioxygenase 1 |
| IDO2 | NA | indoleamine 2,3-dioxygenase 2 |
| TDO2 | 3.97 | tryptophan 2,3-dioxygenase |
| KMO | -0.42 | kynurenine 3-monooxygenase |
| KYNU | 2.78 | kynureninase |
| HAAO | -0.03 | 3-hydroxyanthranilate 3,4-dioxygenase |
| QPRT | 0.076 | quinolinate phosphoribosyltransferase |
| NMRK1 | 0.33 | nicotinamide riboside kinase 1 |
| NMRK2 | NA | nicotinamide riboside kinase 2 |
| NMNAT1 | 0.03 | nicotinamide nucleotide adenylyltransferase 1 |
| NMNAT2 | 3.76 | nicotinamide nucleotide adenylyltransferase 2 |
| NMNAT3 | -0.10 | nicotinamide nucleotide adenylyltransferase 3 |
| Naprt | -0.71 | nicotinate phosphoribosyltransferase |
| NAMPT | 0.83 | nicotinamide phosphoribosyltransferase |
| Nadsyn1 | 0.4 | NAD synthetase 1 |
| SIRT1 | -0.16 | sirtuin 1 |
| SIRT2 | 0.30 | sirtuin 2 |
| SIRT3 | -0.61 | sirtuin 3 |
| SIRT4 | -0.012 | sirtuin 4 |
| SIRT5 | -0.23 | sirtuin 5 |
| SIRT6 | -0.11 | sirtuin 6 |
| SIRT7 | 0.27 | sirtuin 7 |
| CD38 | -1.71 | CD38 molecule |
| PARP1 | -0.20 | poly(ADP-ribose) polymerase 1 |
| PARP2 | -0.68 | poly(ADP-ribose) polymerase 2 |
| PARP3 | 0.04 | poly(ADP-ribose) polymerase 3 |
| PARP4 | 0.48 | poly(ADP-ribose) polymerase 4 |
| PARP6 | -0.33 | poly(ADP-ribose) polymerase 6 |
| PARP8 | 0.22 | poly(ADP-ribose) polymerase 8 |
| PARP9 | 1.64 | poly(ADP-ribose) polymerase 9 |
| PARP10 | 1.1 | poly(ADP-ribose) polymerase 10 |
| PARP11 | NA | poly(ADP-ribose) polymerase 11 |
| PARP12 | 1.14 | poly(ADP-ribose) polymerase 12 |
| PARP14 | 1.22 | poly(ADP-ribose) polymerase 13 |
| PARP15 | 0.96 | poly(ADP-ribose) polymerase 15 |
| PARP16 | -0.49 | poly(ADP-ribose) polymerase 16 |
| ART1 | 4.06 | ADP-ribosyltransferase 1 |
| ART3 | 1.24 | ADP-ribosyltransferase 3 |
| ART4 | 1.29 | ADP-ribosyltransferase 4 |
| ART5 | 3.73 | ADP-ribosyltransferase 5 |

### 3.6 AKR1C inhibitors could bind with AKR1C3 at different locations, as predicted by molecular docking stimulations

Docking simulations with an NADP+-bound AKR1C3 model (Fig. 4A–C) showed that the highest affinity docking inhibitor was MPA, with the lowest binding free energy (ΔG = −11.54 kcal/mol). MCFLA showed the second highest affinity, with ΔG = −7.06 kcal/mol, and MLS showed the lowest affinity (−6.47 kcal/mol). These results are consistent with the IC50 values of the inhibitors on AKR1C3, with MPA having the lowest and MLS having the highest IC50 value (Fig. 2A).

**Fig. 4.**
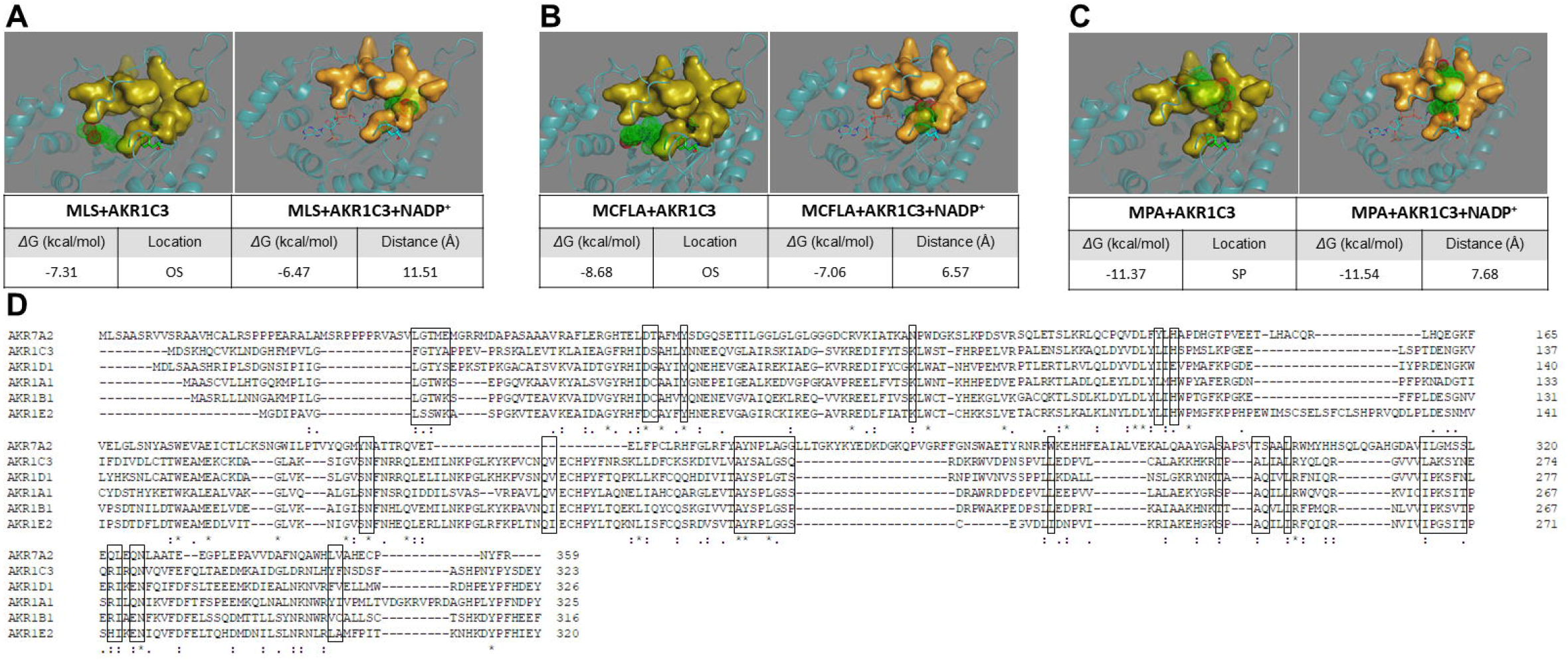
Molecular docking simulation results of AKR1C3 and three inhibitors. A, Meclofenamic acid (MCFLA) with AKR1C3 alone or AKR1C3 and NADP+. B, Methyliasmonate (MLS) with AKR1C3 alone or AKR1C3 and NADP+. C, Medroxyprogesterone acetate (MPA) with AKR1C3 alone or AKR1C3 and NADP+. OS, oxygen site; SP, steroid-binding pocket. Distance was measured between the nitrogen atom of the nicotinamide ring and the carbonyl oxygen of the inhibitor. AKR1C3 is represented in cartoon style. NADP+ is shown as a stick model, and Tyr-55 and His-117 are shown as a stick-ball model. NADP+, Tyr-55, and His-117 contain an oxygen site (OS). The steroid pockets (SPs) are displayed in surface style. D, The alignment of representative members of the AKR enzyme family (AKR1A1, AKR1B1, AKR1C3, AKR1D1, AKR1E2, and AKR7A2). The residues involved in the NADP+ binding are placed in the box.

If NADP+ was removed from the AKR1C3 structure and only inhibitors were docked with AKR1C3 apoenzyme, the order did not change, but we could see considerable differences in their inhibitor binding location (Fig. 4A-C). MCFLA and MLS occupied the oxygen site composed by Tyr-55, His-117, and NADP+, while MPA was tightly bound to the steroid-binding site. In the NADP+-bound AKR1C3 model, the distance between the nitrogen atom of the nicotinamide ring of NADP+ and the carbonyl oxygen of the inhibitors was consistent with the notion that MCFLA was closer to NADP+ (~6.57 Å) than MPA (~7.68 Å). These results indicate that MPA is the best selective inhibitor of AKR1C3, while MCFLA partially interacts with the NADP+ binding site, showing a relatively poor selectivity.

## 4 Discussion

It has been reported that AKR1C enzymes are overexpressed in several cancer cell types, such as bladder cancer cells and gastric cancer cells, contributing to the resistance to Cis-platin treatment[2–6,11]. Some cytotoxic lipid peroxidative products have been mentioned as targets of AKR1Cs that are involved in Cis-platin resistance[6]. However, the role of AKR1Cs in the mechanism underlying Cis-platin resistance remains unclear. We found that AKR1C inhibitors could reverse resistance of HepG2-RC cells, even though AKR1Cs were not significantly upregulated in HepG2-RC cells. The effects of siAKR1Cs on HepG2-RC cells were comparable to those of the inhibitor MPA, but could only partially explain the effects of MCFLA. This fact could be explained by that MPA is a steroid analog, which shows a high selectivity, while MCFLA inhibits not only AKR1C enzymes but also other AKR enzymes. MCFLA belongs to the class of non-steroid anti-inflammatory inhibitors, which have been reported to also inhibit cyclooxygenases besides AKR1Cs, indicating its poor selectivity[12]. As shown in Fig. 4D, all AKR enzymes share a common set of residues to bind NADP(H). As shown in Fig. 4B, MCFLA preferred the oxygen site rather than the steroid pocket, indicating that MCFLA inhibits AKR1C3 by disturbing NAD(P) (H) binding.

These results indicate that MCFLA could inhibit some other AKR enzymes besides AKR1Cs. This could explain why MCFLA causes a stronger reversal of Cis-platin resistance of HepG2-RC cells than MPA. It is likely that NAD(P)H plays a key role here because it is a reducing force and a product of NAD(P)H-dependent oxidoreductases. Reprogramming energy metabolism is considered as a hallmark of cancer cells in which NAD(P) or NAD(P)H levels are increased[13–15]. It has been reported that some NAD(P)H-dependent oxidoreductases, such as ALDHs, increase NAD(P)H levels in the cytosol of cancer cells, which then serves as an electron source[16].

We also found that the amount of total NAD (NAD++NADH) in both HepG2 and HepG2-RC cells was approximately four-fold higher than in HL-7702 cells. Moreover, the ratio of NADH/NAD+ in HepG2-RC cells was increased as much as seven-fold compared to HepG2 cells. The increased ratio could be explained by two steps (Fig. 5). First, NADH is produced continuously in hepatic cancer cells. This is in contrast to the situation in normal cells, where AKRs function as reductases[1]. However, the continuously biosynthesized NAD in hepatic cancer cells would make NAD(P)H-dependent enzymes catalyzing reactions in one direction, from NAD(P) to NAD(P)H. Second, the produced NAD(P)H would deal with Cis-platin-induced peroxidative products directly or indirectly. Usually, NADPH maintains glutathione at the reduced state, which could detoxify reactive oxygen species. However, this could not alter the NAD(P)H/NAD(P) ratio too much, because each reaction is reversible. It has also been reported that no alteration of resistance to Cis-platin or oxaliplatin occurred after GSH depletion in oxaliplatin-resistant human gastric adenocarcinoma TSGH cells[17], indicating GSH is not involved in the Cis-platin resistance.

**Fig. 5.**
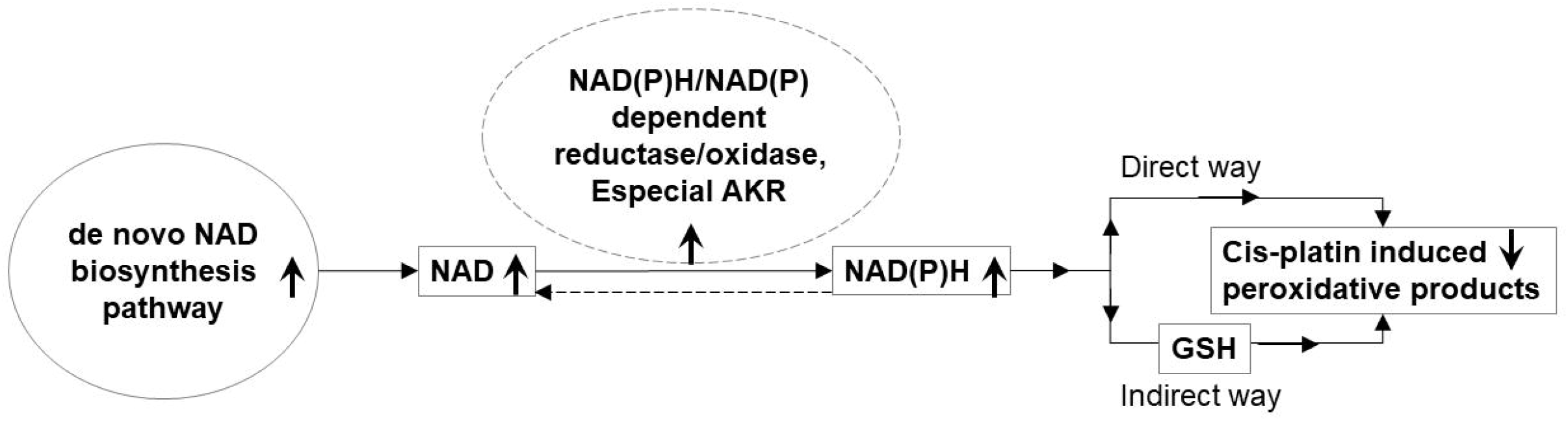
A possible role of NAD(P)H/NAD(P)-dependent reductase/oxidase in the mechanism underlying Cis-platin resistance. Arrows pointing upwards indicate upregulation in HepG2-RC cells; arrows pointing downwards indicate downregulation in HepG2-RC cells.

Besides GSH, NAD(P)H could also react rapidly with moderately oxidizing radicals to repair biomolecules[18]. So, NAD(P)H could also work as a directly operating antioxidant that scavenges radicals as NAD(P)H* forms. These forms would exist in the cell for a relatively long time and keep the intracellular concentration of free NAD(P)H very low. Therefore, NAD(P)H-dependent enzymes could catalyze NAD(P) to NAD(P)H continuously, resulting in a high ratio of NAD(P)H/NAD(P). If these enzymes were widely inhibited by a poor selective inhibitor, such as MCFLA, NADH would not accumulate anymore, and Cis-platin resistance would be suppressed as well.

## 5 Conclusions

In summary, it is believed that NAD(P)H-dependent oxidoreductases, especially AKRs, produce NADH in HepG2 cells to overcome Cis-platin-induced cytotoxicity. According to this notion, chemotherapy with inhibitors, which could compete with NAD(P) in most oxidoreductases, could lead to a better reversal of Cis-platin resistance in Cis-platin-resistant cancer cells.

## 7 Author Contributions

Yuanhua Yu conceived-designed experiment. Tingting Sun and Le Gao analyzed data and wrote and edited the main manuscript text. Xin Wang, Xue Sun and Rui Guo performed in vitro experiments. All authors read and approval of the manuscript.

## 9 Acknowledgments

We thank LetPub (www.letpub.com) for its linguistic assistance during the preparation of this manuscript.

## 10 Supplementary Material

Additional tables(Table.S1., Table.S2., Table.S3.)

