## Supplemental Data 1 for "Analysis of the mechanism of Aldo-keto reductase dependent cis-platin resistance in HepG2 based on transcriptomic and NADH metabolic analysis"

Table.S1. The sequences of oligo nucleic acid used in qRT-PCR and Knock-down experiment.

| Name | Sequence | Usage |
| --- | --- | --- |
| AKR1C3-Fw  AKR1C3-Rv  siAKR1Cs  siNC | CATTGGGGTGTCAAACTTCA  CCGGTTGAAATACGGATGAC  AACACCUGCACGUUCUGUCUGAUGC  UUCUCCGAACGUGUCACGUTT | For qRT-PCR  For qRT-PCR  For knock-down  For knock-down |

Table.S2. TOP 10 of upregulated genes in HepG2-RC compared to HepG2

| Gene | log2(HepG2-RC/HepG2) | Annotation |
| --- | --- | --- |
| SPINK6  TMEM140  CLEC3A  PSG2  SLC6A15  CLIC5  CASP14  CDH10  BRINP3  DSC2 | 11.71  9.87  8.93  8.81  8.24  8.13  8.05  7.89  7.77  7.63 | serine peptidase inhibitor Kazal type 6  transmembrane protein 140  C-type lectin domain family 3 member A  pregnancy specific beta-1-glycoprotein 2  solute carrier family 6 member 15  chloride intracellular channel 5  caspase 14  cadherin 10  BMP/retinoic acid inducible neural specific 3  desmocollin 2 |

Table.S3. TOP 10 of downregulated genes in HepG2-RC compared to HepG2.

| Gene | log2(HPEG2-RC/HPEG2) | Annotation |
| --- | --- | --- |
| POTEJ  SPDEF  TCP10L  ARX  PPP1R1B  LOC107986354  AGR2    CYP4Z1  BPIFA2  NLRP9 | -5.92  -5.28  -4.92  -4.78  -4.67  -4.55  -4.52    -4.49  -4.42  -4.35 | POTE ankyrin domain family member J  SAM pointed domain containing ETS transcription factor  t-complex 10 like  aristaless related homeobox  protein phosphatase 1 regulatory inhibitor subunit 1B  uncharacterized  anterior gradient 2, protein disulphide isomerase family member  cytochrome P450 family 4 subfamily Z member 1  BPI fold containing family A member 2  NLR family pyrin domain containing 9 |
